# Are microbes growing on flowers evil? Effects of old flower microbes on fruit set in a wild ginger with one-day flowers, *Alpinia japonica* (Zingiberaceae)

**DOI:** 10.1101/2021.06.28.450259

**Authors:** Nuria Jiménez Elvira, Masayuki Ushio, Shoko Sakai

## Abstract

Flowers are colonized and inhabited by diverse microbes. Plants rapidly replace flowers with short lifespan, and old flowers senesce. This may contribute to avoiding adverse effects of the microbes. In this study, we investigate if the flower microbial community on old flowers impedes fruit and seed production in a wild ginger with one-day flowers. We inoculated newly opened flowers with old flower microbes, and monitored the effects on fruit and seed set. We also assessed prokaryotic communities on the flowers using amplicon sequencing. We found six bacterial amplicon sequence variants (ASVs) whose proportions were increased on the inoculated flowers. These ASVs were also found on flower buds and flowers that were bagged by net or paper during anthesis, suggesting that they had been present in small numbers prior to flowering. Fruit set was negatively associated with the proportions of these ASVs, while seed set was not. The results suggest that old flowers harbor microbial communities different from those at anthesis, and that the microbes abundant on old flowers negatively affect plant reproduction. Though the short lifespan of flowers has gotten little attention, it might be an essential defense mechanism to cope with antagonistic microbes that rapidly proliferate on the flowers.

## 1. INTRODUCTON

Flowers are an elaborate device to exchange pollen among conspecific individuals. Instead of moving around to find a potential mate, the plants disperse and receive pollen relying on different vectors such as insects and wind to achieve sexual reproduction. As a consequence, flowers are also exposed to microbial colonization under natural conditions (Burdon et al. 2018; Kaltz and Shykoff 2001; Ushio et al. 2015). Flowers have various habitats for microbes. Flowers attract pollinators by secreting nectar, which is rich in sugars and often contains other nutrients, such as amino acids and lipids (Roy et al. 2017). The stigma is a germination bed for pollen grains connected to a growth chamber for pollen tubes. It maintains humidity and nutrients necessary for pollen tube growth (Taylor and Hepler 1997), which would also be beneficial for microbes. Recent studies using high-throughput sequencing have revealed highly diverse microbial communities on flowers of many species in a wide range of habitats (Gaube et al. 2021; Massoni et al. 2020; Wei and Ashman 2018).

Today, flower microbes are increasingly recognized as essential components in the ecology and evolution of plant reproduction (reviewed in Rebolleda-Gómez et al. 2019; Vannette 2020). Not surprisingly, some flower microbes are known to have strong negative effects on the survival and reproduction of the host plant. Many pathogens use flowers, especially the stigma and nectary, as a point of entry. Their infection of flowers often results in fruit abortion and systemic infection (Bubán et al. 2003; Vanneste 2000). Other microbes impede various processes of pollination and fertilization. Nectar yeasts are reported to modify pollinator foraging patterns and reduce pollination success (Herrera et al. 2013). *Acinetobacter* bacteria that commonly inhabit floral nectar exploit pollen nutrition by inducing pollen germination and bursting (Christensen et al. 2021). Most studies of these phenomena examined the effects of an individual microbial species or of separate strains rather than that of the whole community.

In general, even newly opened flowers already have microbes at least on some flower tissues. Their abundance increases over time on individual flowers (Vannette 2020). Flowers often have characteristics that may suppress microbial growth, such as flower volatiles (Burdon et al. 2018; Huang et al. 2012), reactive oxygen (McInnis et al. 2006) and secondary compounds (Dobson and Bergstrom 2000; Palmer-Young et al. 2019). An extremely short lifespan (from one to several days) of flowers (Primack 1985), especially in hot habitats (Bawa 1990), is also thought to reduce the chance of microbial infection and the proliferation of microbes (Kaltz and Shykoff 2001; Rogers 2006; Shykoff et al. 1996). Unlike the reproductive structures of mammals, flowers, the reproductive structures of the plants, are only retained while they are needed, and are developed *de novo* every season in perennial species (Rogers 2006). We do not know yet, however, to what extent microbes that are suppressed by abscission of flowers are potentially detrimental to plant fecundity.

In this study, we examine if the flower microbial community on old flowers impedes fruit and seed production in a wild ginger, *Alpinia japonica* (Zingiberaceae), that produces short-lifespan flowers. This perennial herb grows on the humid forest floor of evergreen and deciduous forests in western Japan. It flowers in June, when the climate is hot and humid. Its flowers open in the morning and wilt at sunset within the same day. Therefore, its flowering coincides with the optimal season for microbial growth, while it may avoid the negative effects of microbes by renewing flowers daily. To imitate microbial communities that would appear on the flowers of *A. japonica* if their longevity were longer, we inoculate microbes from old flowers onto newly opened flowers in the morning. Firstly, we characterize the prokaryote communities on *A. japonica* flowers. Secondly, we identify prokaryotes that significantly increased after inoculation by comparing prokaryotic communities between the inoculated flowers and open flower controls. In addition, we examine whether these prokaryotes colonize via air or via flower visitors after flowering, or if they had already been present before anthesis. We covered the flowers by net or paper bags to exclude microbial colonization by flower visitors or by both flower visitors and air, respectively. If the old-flower microbes colonize only after anthesis, bagging would reduce the occurrences of these microbes. Lastly, we observe fruit and seed sets of the flowers under the treatments. We expected that the flowers that had more old-flower microbes would show lower fruit and seed set if microbe proliferation on flowers is generally to be avoided.

## 2. MATERIALS AND METHODS

### 2.1 Study species

*Alpinia japonica* (Zingiberaceae) is a wild ginger distributed in temperate and subtropical regions of eastern Asia. It is a perennial herb 0.5–0.7 m in height occurring mostly in broad-leaved evergreen forests. Inflorescences are compound racemose with 10–60 flowers (37 on average in this population). Flowers are zygomorphic with a prominent labellum 1.1–1.2 cm in length, and white with red stripes at the margin of the labellum (Fig. 1a). Flowering occurs almost synchronously within a population and lasts for 10–14 days for each plant between late May and June. From one to five flowers open per inflorescence. At anthesis around 0700, the dorsal and lateral petals open, and the labellum inside expands (Fig. 1b). The flowers are known to have volatiles, including 1, 8-cineole, with antimicrobial activities (Asakawa et al. 2017; van Vuuren and Viljoen 2007). The corolla wilts at sunset within the day. The ovary matures into an ellipsoid, three-loculate fleshy capsule ~2 months after flowering. The species is self-compatible, and pollinated by bees, mainly carpenter bees (*Xylocopa appendiculata*) and bumble bees (*Bombus ardens*) (J. Nuria and S. Sakai, unpublished data).

**FIGURE 1.**
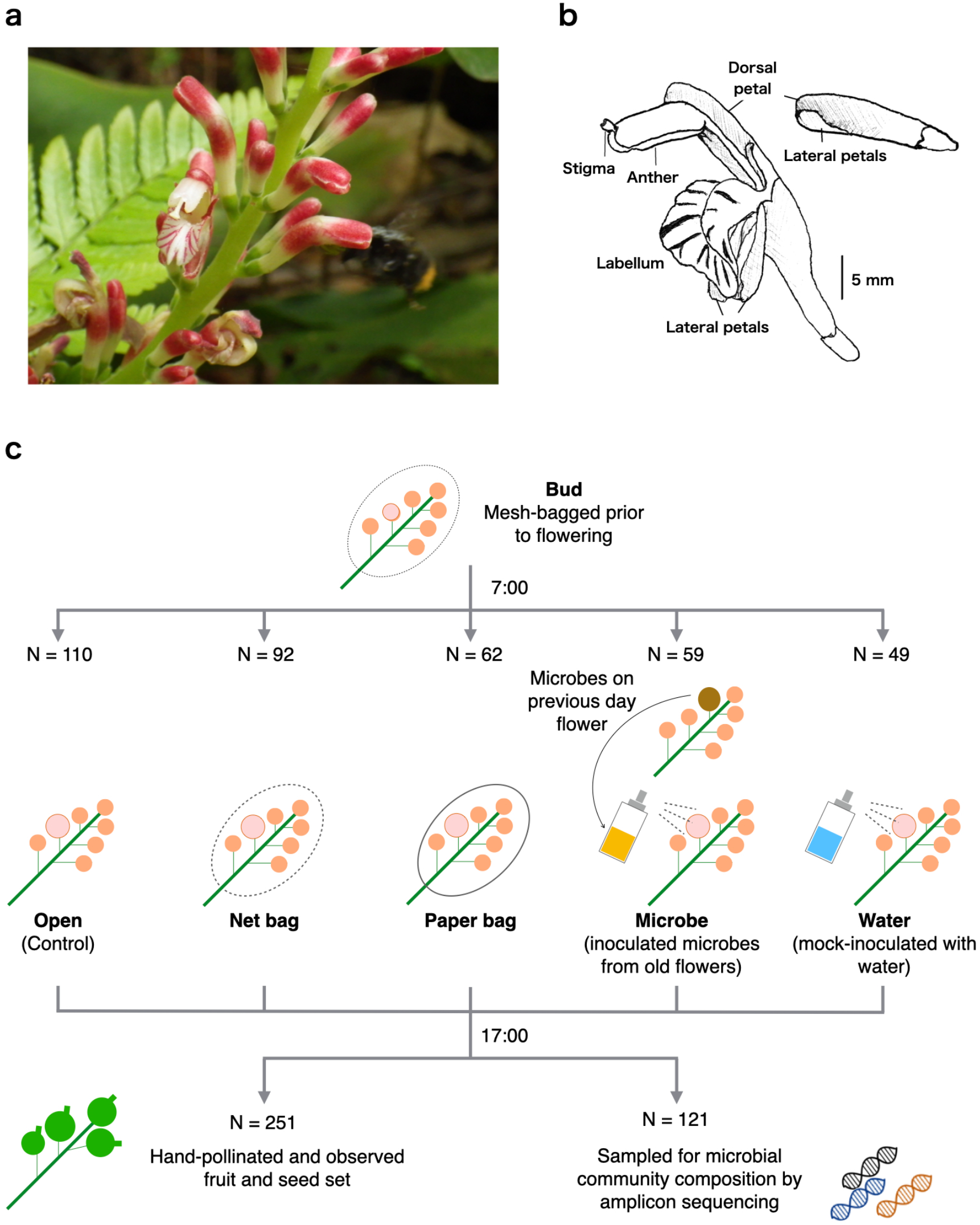
Experimental design used in this study. a, An open flower and flower buds on an inflorescence of *Alpinia japonica*. b, Floral parts of *Alpinia japonica* in open state (left) and bud state (right). c, Experimental and sampling procedures.

### 2.2 Study site

Studies were conducted in May to July of 2018, in Seta Park, Otsu, Shiga Prefecture, Japan (34°50’N, 135°50’E). Seta Park has a young secondary deciduous forest where streams are abundant, which provides an optimal habitat for *A. japonica*. The annual mean temperature is 14.9°C, and annual total rainfall is ~1530 mm in Otsu city (Japan Meteorological Agency, https://www.data.jma.go.jp/). Mean daily maximum temperatures in May, June and July 2018 were 24.0°C, 27.3°C and 33.7°C, respectively.

### 2.3 Flower treatments

*Alpinia japonica* typically presents 2–12 inflorescences per plant at the study site. We selected six plants with at least eight inflorescences with more than 10 flowers on each. Unfortunately, one of them had been damaged, probably by park visitors, before the experiments and sampling. Therefore, we used the remaining five plants (Plant ID = N1, N2, N4, N5, N6). In each target plant, eight inflorescences were tagged and covered with a fine net (Cloth Cabin, Suminoe Teijin Techno, Osaka, Japan) prior to flowering.

We applied one of five treatments to part of the flowers that opened on the inflorescences between 22nd May and 7th June. The flowers under these treatments were used for either identification of microbes on the flowers or for observation of fruit set (Fig. 1c). We could not use the same flower for both, because we should collect the flower on the same day for the former, while we should leave the flower intact for the latter. The five treatments were as follows: 1) open control (OT), in which the mesh bag on the inflorescence was removed during 0700–1700, so that microbes were transmitted to the flower by both air and pollinators; 2) insect exclusion by a net bag (NB), in which the inflorescence was left covered with the net, which allowed air but not insects to get through; 3) paper bagged (PB), in which the mesh bag covering the inflorescence was replaced by a new paper bag (Grape Bag, DAIICHI VINYL, Fukui, Japan) during 0700–1700 to exclude air flow and insect visits; 4) open and microbial inoculation (MI) and 5) open and water sprayed mock (WI) treatments. In MI, the mesh bag on the inflorescence was removed as in OT, and the flower was sprayed with microbial solution. During the spray inoculation, the focal flower was isolated from the rest of the inflorescence by a funnel-shaped plastic sheet to avoid contamination. The microbial suspension for the inoculation was prepared by soaking old flowers in distilled water. Nine old flowers that had opened the previous day from 1–3 individuals were collected from individuals that did not receive the treatments. Microbes were detached by ultrasonic dispersion in 10 ml of distilled water and kept at 4°C until the inoculation (maximum of 36 hours). Inoculation was conducted on eight different days, and the same inoculant was used for all flowers under MI treatment on the same day. In WI, the mesh bag covering the inflorescence was removed as in OT, and the flower was sprayed with distilled water as a control for the physical effect of the MI treatment. We did not conduct the treatments on cloudy or rainy days. Flowers that opened on the same day on the same inflorescence were assigned to the same treatments in the case of NB and PB, while OT, MI and WI were occasionally mixed on an inflorescence. This is because we bagged the entire inflorescence rather than a single flower to avoid damaging the flowers. The sample size of each treatment is summarized in Table S1.

### 2.4 Identification of microbial communities

To identify microbial communities, we sampled 4–5 flowers with each treatment from each plant. The flowers were cropped at ~1700 of the day of flower opening and separately put into a 5 ml plastic tube after removing the stigma (to use for another study) and petals (Fig. 1b). In addition, five flower buds were collected from each individual to identify microbes that were present inside prior to anthesis with the same procedure. The tube was kept on ice until being brought to the lab within 4 hours after sampling and kept at –20°C until further processing.

To detach microbes from the flower surface, we added 3 ml of phosphate-buffered saline solution (PBS buffer, pH 7.4, Nippon Gene Co., Ltd.) to the tube and sonicated it using an Ultrasonic Disruptor Handy Sonic UR-21P (Digital Biology) at Level 5 power for 30 sec. Microbes in the PBS solution were then filtered using a Sterivex Millipore filter (Sterivex-HV 0.22, Merck). Residual PBS on the Sterivex filter was removed using a Vac-Man® Laboratory Vacuum Manifold (Promega).

Microbial DNA was extracted with DNeasy Blood & Tissue Kits (QIAGEN) following the manufacturer’s instructions. A portion of the 16S small-subunit ribosomal gene was PCR-amplified using the primer pairs 515F (5’-GTG YCA GCM GCC GCG GTA A-3’) and 806R (5’-GGA CTA CNV GGG TWT CTA AT-3’) (Caporaso et al. 2011) with Nextera DNA index adapters. First PCR was performed in 12 μl reactions, each containing 1 μl of DNA template, 6 μl of KAPA HiFi HotMasterMix (Kapa Biosystems) and 0.7 μl of each primer (5 μM). The first PCR was conducted with a temperature profile of 95°C for 3 min, followed by 35 cycles at 98°C for 20 s, 60°C for 15 s, 72°C for 30 s, and a final extension at 72°C for 5 min, followed by storage at 4°C until use. Triplicate PCR products were pooled and purified with Agencourt AMPure XP (Beckman Coulter). P5/P7 Illumina adaptors were then added in a second PCR using fusion primers with 8-mer index sequences for sample identification (Hamady et al. 2008) (forward, 5’-AAT GAT ACG GCG ACC ACC GAG ATC TAC AC-[8-mer tag]-TCG TCG GCA GCG TCA GAT GTG TAT AAG AGA CAG −3’; reverse, 5’-CAA GCA GAA GAC GGC ATA CGA GAT-[8-mer tag]-GTC TCG TGG GCT CGG AGA TGT GTA TAA GAG ACA G-3’). This preparation had a total volume of 24 μl with double the amount of each reagent in the first PCR and 10-times diluted first PCR products as DNA template. The temperature profile was 95°C for 3 min, followed by 12 cycles at 98°C for 20 s, 68°C for 15 s, 72°C for 15 s, a final extension at 72°C for 5 min, and 4°C until use. Second PCR products were pooled at equal volumes after a purification/equalization process with the same AMPure XP protocol and E-Gel SizeSelect II Agarose Gel kit (Invitrogen). DNA quantitation was performed using a Qubit® dsDNA HS Assay Kit (Thermo Fisher Scientific Inc). The pooled library was sequenced on an Illumina Miseq using Reagent Kit V2 (500 cycle Nano Kit).

We processed the sequence data following Ushio (Ushio 2019). The raw MiSeq data were converted into FASTQ files using the bcl2fastq program provided by Illumina (bcl2fastq v2.18). The FASTQ files were then demultiplexed using the command implemented in Claident (http://www.claident.org; (Tanabe and Toju 2013)). Demultiplexed FASTQ files were analysed using the ASV method implemented in the package DADA2 (Callahan et al. 2016) of R (R Core Team 2016). At the quality filtering process, forward and reverse sequences were trimmed at the length of 215 and 160, respectively, using DADA2::filterAndTrim() function.

Taxonomic identification was performed for ASVs (Amplicon Sequence Variants; proxy for Operational Taxonomic Units) inferred using DADA2 based on the query‐centric auto‐k‐nearest‐neighbour (QCauto) method (Tanabe and Toju 2013) and subsequent taxonomic assignment with the lowest common ancestor algorithm (Huson et al. 2007) using the “merge” database and clidentseq and classigntax commands implemented in claident v0.2.2018.05.29. The reads identified as Bacteria or Archaea were used for further analysis. Taxonomy, ASV table, and sample information were combined into a single R object for downstream manipulation using the R package phyloseq (McMurdie and Holmes 2013). We also removed ASVs that only occurred in negative control samples. Furthermore, potential contaminant ASVs were identified statistically based on prevalence in negative control samples using the R package decontam (Davis et al. 2018) based on ASV prevalence and a threshold of 0.5. This resulted in elimination of 87 of 2915 16S ASVs.

### 2.5 Fruit set and seed production

The treated flowers for the observation of fruit set under all the treatments were hand-pollinated around 1700 using outcrossing pollen to ensure that the fruit set was not limited by pollen. Pollen was taken with a wood toothpick from a cut flower from a pollen donor plant, of which inflorescences had been covered with a mesh bag prior to anthesis to prevent removal of pollen by flower visitors. In total, 251 pollinated flowers were monitored for fruit and seed set (Table S1). Six weeks after the treatment, we surveyed whether the fruits under the treatments were still retained on the inflorescence, and collected the fruits (131 fruits) and counted developing seeds and aborted ovules inside. At that time, the fruits were still green but fully plump.

### 2.6 Statistical analysis of microbial community data

To examine differences in the prokaryotic diversity among the treatments, we evaluated the Shannon diversity index of each sample using the phyloseq::estimate_richness() function. Significant differences in the index among treatments were tested using Generalized Linear Mixed Model (GLMM). Prior to the analysis, we confirmed that distributions of the index were not significantly deviated from the normal distribution (Shapiro-Wilk test, W = 0.983, p = 0.0759). We constructed a GLMM with the treatment as a fixed term and Plant ID as a random factor using the R package lme4 (Bates et al. 2015). Significance of the fixed term was tested by comparing the full model and the model without the fixed term using the anova() function.

We examined variation in the structure of the prokaryotic communities using the ASVs that were present in 20 or more flower samples (36 ASVs satisfied this criterion). The frequency of each ASV was divided by the number of the total sequences of the 36 ASVs in each sample to convert it to a proportion.

To compare microbial communities among flowers under the different treatments, we calculated the Bray–Curtis dissimilarity index using the package vegan (Oksanen et al. 2010) in R. We performed a permutational multivariate analysis of variance (PERMANOVA) using the vegan::adonis function to examine if flowers of the same individuals nested under the same treatment have similar microbial communities. Ordinations were plotted with nonmetric multidimensional scaling (NMDS) using the vegan::metaMDS procedure. We then calculated the distances of each flower sample from the centroid of the treatment. Variation in the distances among the treatments were tested using vegan::anova for betadisper object.

To find ASVs that tend to co-occur in the samples, we conducted hierarchical clustering of ASVs based on Bray–Curtis dissimilarity using Ward’s minimum variance algorithm using the hclust procedure. The algorithm merges clusters that minimize the increase in the sum of squared distances from the cluster centroid. We chose the number of clusters that maximize the silhouette index (Rousseeuw 1987). The index was calculated using silhouette in the R package cluster (Maechler et al. 2001). The proportions of the ASVs in the sample were visualized by a heatmap created using the R package ComplexHeatmap (Gu et al. 2016). We identified the cluster that showed the largest difference in the median proportion in the samples under the microbial inoculate (MI) treatment compared with that under the water sprayed (WI) treatment. Differences in the proportions of the cluster between the MI and WI treatments were also evaluated by the statistic of the Wilcoxon rank sum test. The identification of the ASVs in the cluster was searched against the Nucleotide database of the National Center for Biotechnology Information (NCBI) with BLAST (Altschul et al. 1990). Top matches of the sequences were referred to in order to assign taxonomic identities (Class, Family, Genus) of the ASVs.

### 2.7 Statistical analysis of fruit set and seed set data

To assess the factors associated with fruit set and seed production, we used generalized linear mixed models (GLMMs). To evaluate differences in fruit set among the treatments after controlling for the difference among the individuals, we modeled fruit set (retained/aborted) with the treatments as a fixed factor and the individuals as a random factor using a binomial error distribution. Then we calculated and plotted estimated marginal means of fruit set for the treatments using the ggpredict() function of the R package ggeffects (Lüdecke 2018). Then, to test the association of fruit set with microbial communities, we constructed the second model for fruit set. The model included microbial compositions as a fixed factor and the individuals as a random factor using a binomial error distribution. We identified Cluster 6 to include ASVs that showed the largest increase after the microbial inoculation (see below). Therefore, we used the average proportion of Cluster 6 of each treatment as the variable for the microbial composition. We evaluated the significance of the fixed factor by the Wald test.

We also modeled seed set (retained/aborted) and tested its association with microbial composition. The model included the proportion of Cluster 6 as a fixed factor and the individuals and fruit as nested random factors. We fitted the model with a binominal error distribution. The GLMM analyses for fruit and seed sets were performed using the R package lme4 (Bates et al. 2015).

## 3. RESULTS

### 3.1 Microbial communities

We obtained 839,113 reads of prokaryotic sequences, which were clustered into 2,828 ASVs. We have removed samples that had < 250 prokaryotic reads (12 samples). This reduced the number of samples to 134. According to the rarefaction curves (Fig. S1), the sequencing captured most prokaryotic diversity in these 134 samples. The median number of reads in the remaining samples was 1354.5. The microbial communities on *Alpinia japonica* flowers were by far dominated by the bacterial phylum Proteobacteria, which accounted for 47.8 % in the samples on average (Fig. 2). At the class level, Gammaproteobaceteria was the most dominant (16.5 %), followed by Alphaproteobacteria (6.1 %) and Betaproteobacteria (5.8 %). The Shannon diversity index of the samples varied among samples, and the majority fell between 1.5 and 3.5 (Fig. S2). The diversity index was significantly different among the treatments, and was the lowest in the paper bag (PB) treatment (GLMM, χ^2^ = 22.23, p = 0.0005).

**FIGURE 2.**
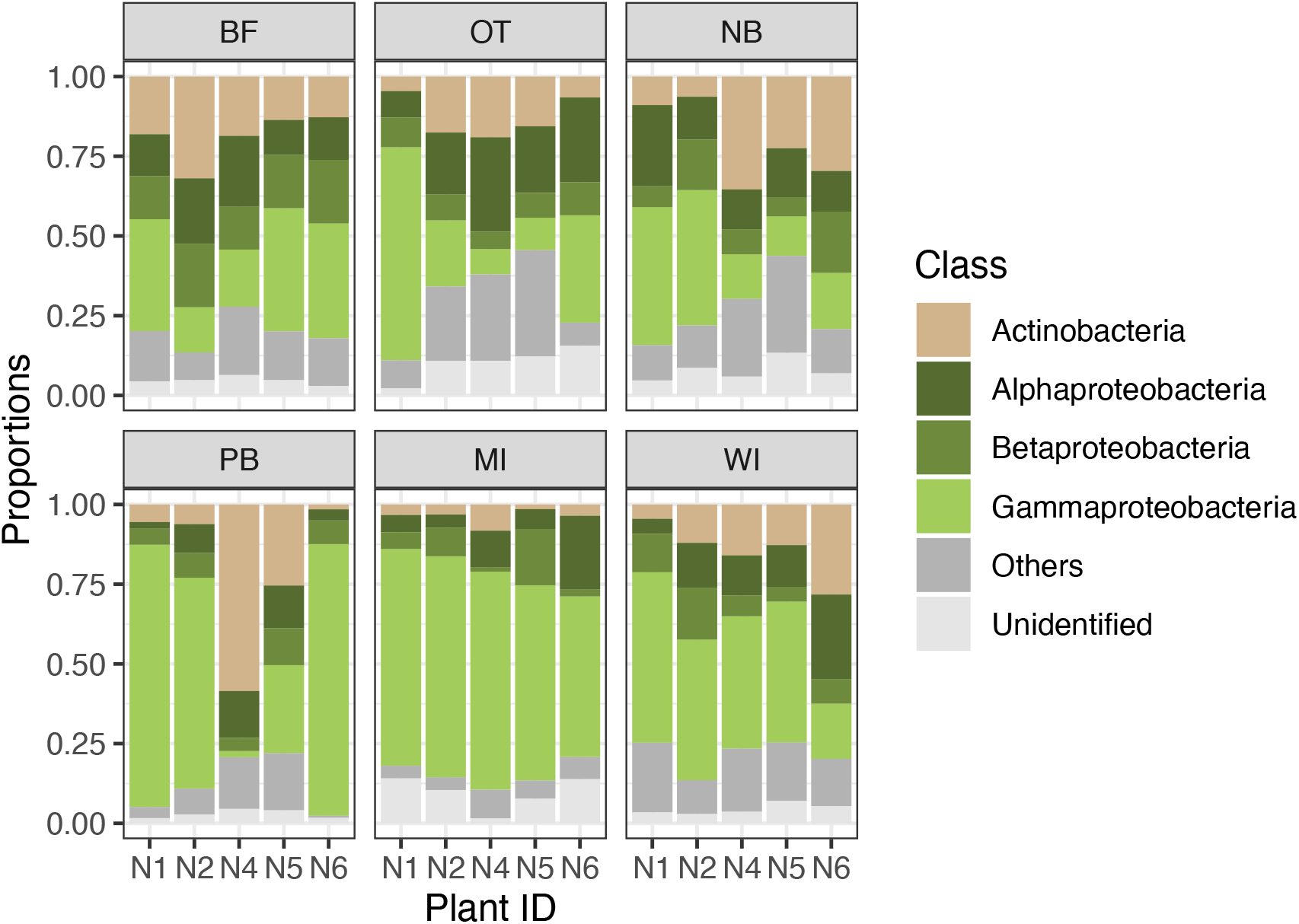
Composition of prokaryotic communities on the flowers of *Alpinia japonica*. Proportions in the samples of the same treatment in the same individuals were averaged. See Table S1 for sample sizes.

The 36 ASVs that appeared in 20 or more samples were identified as bacteria, and accounted for 75.7 % of the prokaryotic sequences. NMDS plots of the microbial communities revealed that the flowers under different treatments in each plant individual had different microbial communities, while the same treatment on different plant individuals did not always result in microbial communities with similar compositions. For example, the OT flowers are plotted on the right of the flower bud samples in the plant N1, while the OT flowers are on the left side of the buds in N4 and N5 (Fig. S3a). Only MI treatment resulted in the conversion of the prokaryotic communities. PERMANOVA indicated that the communities were significantly different among individuals (F = 4.89, R^2^ = 0.100, p < 0.0001) and among the treatments within each individual (F = 2.85, R^2^ = 0.366, p < 0.0001). Variation in the microbial communities within a treatment was significantly different among the treatments (ANOVA, df = 5, F = 8.60, p < 0.0001), and it was the lowest in the MI treatment, as suggested from the NMDS plots (Fig. S3b).

The cluster analyses grouped the 36 ASVs into 11 clusters (Fig. 3, Table S2). The first and second most dominant clusters of the 11 include 4 and 6 ASVs and accounted for 33.0 % and 30.6 % of the sequence reads of the 36 ASVs, respectively (Table S2). Cluster 6, which accounted for the second highest proportion, showed the largest difference in the proportion between the MI and WI treatments, followed by Cluster 2, the most dominant cluster (Table S2, Fig. S4). The Wilcoxon rank-sum test showed that the difference between the treatments was the most significant for Cluster 6 (w = 444, p < 0.0001 for Cluster 6), while Clusters 2 and 5 also showed significant differences (w = 78.0, p = 0.0001 for Cluster 2 and w = 127.0, p = 0.0071 for Cluster 5).

**FIGURE 3.**
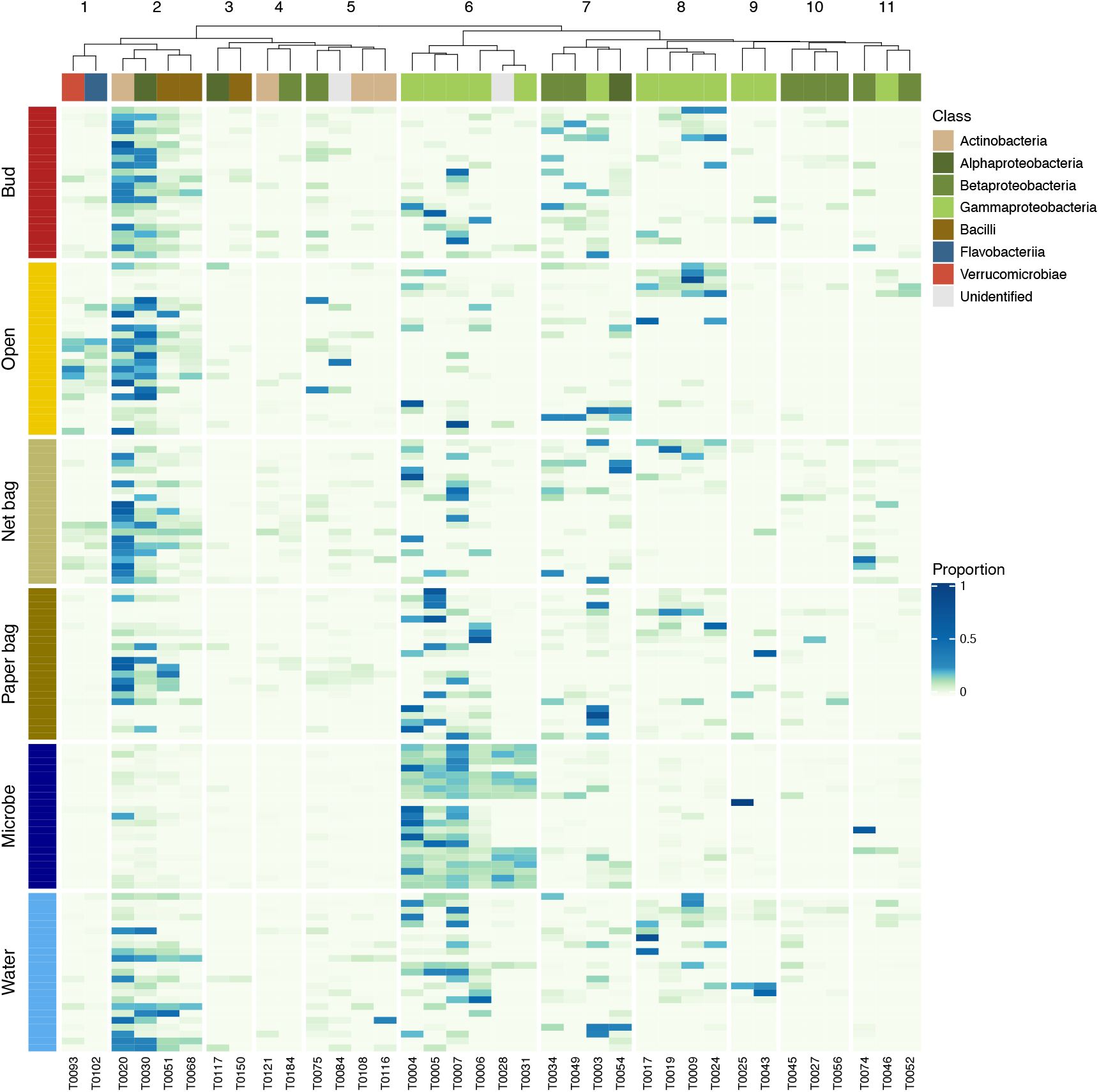
Heatmap of the prokaryotic communities of flower buds and flowers under different treatments. Samples were vertically arranged and grouped by the treatments as shown on the right. ASVs were clustered hierarchically based on Bray–Curtis dissimilarity using Ward’s minimum variance algorithm. The class of each ASV is shown by the color below the tree. The color intensity in each panel shows the proportion in a sample, referring to the color key on the left.

The Claident pipeline identified the most abundant 4 ASVs out of the six in Cluster 6 as belonging to *Pseudomonas* (T00005, T00006 and T0007) and *Erwinia* (T0004). The three *Pseudomonas* ASV sequences were different for five to eight bases among 253 (2.0–3.0 %). Blast search further identified the other two ASVs (T00028 and T00031) in the dataset as *Luteibacter* (Rhodanobacteraceae, Gammaproteobacteria) and *Massilia* (Oxalobacteraceae, Betaproteobacteria) (Table 1).

**TABLE 1.** The 6 ASVs in Cluster 6. Taxonomic information was based on Claident (shown by normal face) supplemented by BLAST search (shown by bold face with percent of identical bases). The results of BLAST search of the ASVs in Cluster 6, and their percent of matches.

| ASV | Phylum | Class | Family | Genus | Percent identity<sup>1</sup> | Average proportion among samples<sup>2</sup> |
| --- | --- | --- | --- | --- | --- | --- |
| T0004 | Proteobacteria | Gammaproteobacteria | **Erwiniaceae** | ***Erwinia*** | 100.0 | 0.086 |
| T0005 | Proteobacteria | Gammaproteobacteria | Pseudomonadaceae | *Pseudomonas* |  | 0.062 |
| T0006 | Proteobacteria | Gammaproteobacteria | Pseudomonadaceae | *Pseudomonas* |  | 0.040 |
| T0007 | Proteobacteria | Gammaproteobacteria | Pseudomonadaceae | *Pseudomonas* |  | 0.086 |
| T0028 | Proteobacteria | **Betaproteobacteria** | **Oxalobacteraceae** | ***Massilia*** | 99.2 | 0.017 |
| T0031 | Proteobacteria | Gammaproteobacteria | Rhodanobacteraceae | ***Luteibacter*** | 99.2 | 0.015 |

Note: 1, The highest percentage of the identical bases that resulted from the BLAST search; it is shown only for the ASVs that were subjected to the BLAST search due to incomplete taxonomic assignments by Claident. 2, Average proportions of the ASV of 36 prevalent ASVs used for the heatmap in Figure 3.

### 3.2 Fruit set and seed production

The probability of fruit set of OT predicted by the first GLMM with the treatment as the fixed factor was 57.3 %, and that of MI was the lowest (34.7 %) (Fig. 3b). Those of the other treatments were between these two (Fig. 3b). On the other hand, the proportion of Cluster 6 showed the opposite patterns; it was the lowest in OT and highest in MI. The GLMM including the proportion of Cluster 6 as the fixed factor indicated that Cluster 6 was significantly associated with low fruit set (GLMM, z = – 2.21, p = 0.0271).

Each fruit that remained on the plants under the treatments had 12–24 ovules, 55.6 ± 20 % of which had developed. In contrast to fruit set, seed set was not significantly associated with the microbial composition (GLMM, z = 0.685, p > 0.05).

## 4. DISCUSSION

### 4.1 Prokaryotic community on *A. japonica* flowers

*Alpinia* flowers harbored diverse prokaryote communities comparable with those on flowers of other plant species reported so far (Berg et al. 2017; Müller et al. 2016; Qian et al. 2021; Wei and Ashman 2018). They are dominated by the phylum Proteobacteria, most of which belonged to Gammaproteobactera and Alphaproteobacteria. *Erwinia* and *Pseudomonas* of the Gammaproteobactera are among the most common constituents of the anthosphere (Vannette 2020).

Though these major taxa were shared among the samples, prokaryotic community compositions were significantly different among these plant individuals. The differences already existed in the buds prior to anthesis. Since flower microbes are often shared by leaves of the same plant (Junker et al. 2011; Massoni et al. 2020; Wei and Ashman 2018), the differences might have originated from vegetative parts of the same individuals. The effects of most treatments were not consistent across the individuals (Fig. S3). Although the paper bag treatment decreased the Shannon diversity index (Fig. S2), we did not find a drastic increase of particular ASVs by this treatment (Fig. 3). Growth of a small number of bacteria species in the bag with high humidity and low UV-irradiance might have reduced the diversity (Hayes et al. 2021). Prokaryotic diversity on open and pollinator-excluded net-bagged flowers was not drastically different from that on the flower buds (Fig. S2). It has been reported that flower visitors disperse microbes among flowers (Ushio et al. 2015), and often increase microbial diversity (Allard et al. 2018). Flower visitors might not have had large effects in our system, however, partly due to the relatively low flower visitor activities during the year of this study (N. E. Jimenez and S. Sakai, personal observation).

### 4.2 Identification of bacteria that were increased by inoculation

We identified a group of ASVs that showed substantial changes after inoculation of old-flower microbes. The cluster analyses grouped the ASVs into 11 clusters based on the co-occurrence among the samples. The microbial inoculation treatment drastically increased Cluster 6 (Figs. 3 and 4), which consists of 6 ASVs. Among the ASVs, four ASVs, one *Erwinia* (T00004) and three *Pseudomonas* (T00005, T00006 and T00007) ASVs accounted for 90 % of the reads of the cluster on average (Table 1). Microbial communities on old flowers might be dominated by these taxa. The bacteria may have already colonized before anthesis, because these bacteria were also found in flower buds at lower abundances as well as in the paper- and net-bagged flowers. The proportions of Cluster 6 were also moderately higher in the bagged and water-sprayed flowers than the proportion under the open control treatment (Fig. S3b). Higher humidity due to these treatments might have enhanced growth of these bacteria. In agriculture, it has often been reported that high humidity or precipitation enhances activities of pathogenic *Erwinia* and *Pseudomonas*, and increases damage by diseases (Billing 1980; Llontop et al. 2020; Pietrarelli et al. 2006).

**FIGURE 4.**
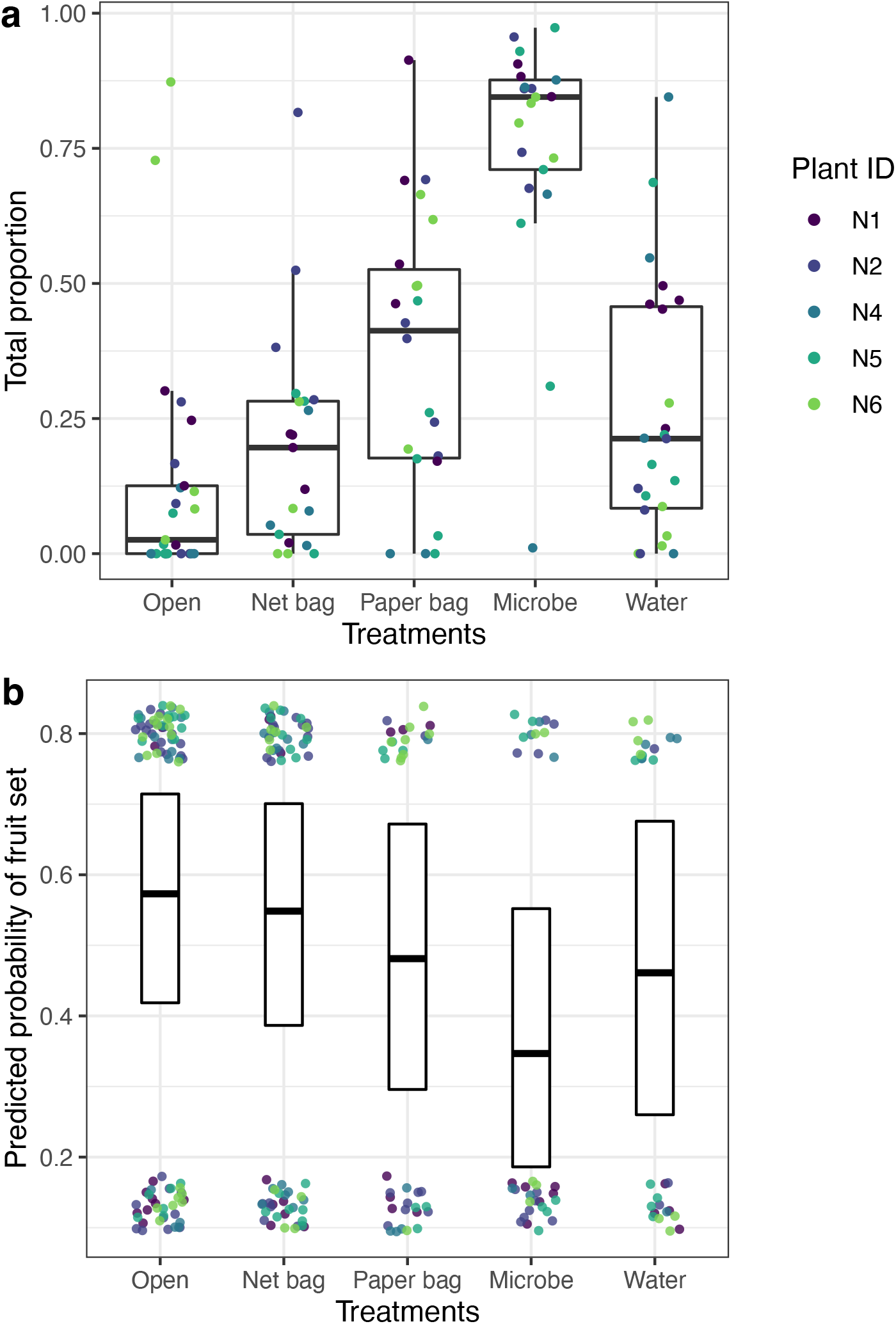
Association between the proportion of Cluster 6 and predicted probability of fruit set. (a) Variation in the total proportions of the bacteria among the treatments. (b) The predicted probabilities of fruit set 6 weeks after flowering under the different treatments by the GLMM with the treatment as the fixed factor. The thick lines and the boxes indicate the predicted values and their 95 % confidential intervals. Distribution of flowers that set fruits (top) or were aborted (bottom) are shown by dots. Colors indicate the five individual plants).

Though the number of studies that investigate microbial communities on flowers is rapidly increasing, no study has examined microbial communities on old flowers as far as we know. Our results suggested that old flowers *A. japonicus* flowers in our field site were dominated by bacteria of *Erwinia* and *Pseudomonas*. Many bacterial species of these genera are frequently found both on flowers and inside the plant and/or on leaves (Vannette 2020). Interestingly, we found all the proteobacterial ASVs in Cluster 6 from flowers of an unrelated dioecious tree, *Mallotus japonicus* (Euphorbiaceae). In particular, T0007 was the most abundant and frequent ASV on male flowers of the plant and was present in 72.3 % of male flower samples (Marre et al. 2021 (T0007 is referred to as ASV02P1in this article), and the dataset used for the study (DRA013108 in DDBJ)). Therefore, we consider that the bacteria of Group 6 may be generalist inhabitants on plants, which proliferate on senesced plant parts. It is not uncommon that flowers of unrelated plant species in the same habitats share flower bacteria species (Massoni et al. 2020; Qian et al. 2021). It should be noted, however, that ASVs in this study are distinguished based on relatively short DNA sequences of 253 base pairs, as is often the case in most anthosphere and phylosphere studies. Besides, we have analyzed the anthosphere of only two plant species so far. To confirm the generalist habit of the bacteria, more detailed examination such as sequencing of additional regions and genes, more extensive sampling and culture isolation would be necessary.

### 4.3 Effects of old-flower microbes on fruit and seed production

Do the microbes that become dominant on old flowers negatively affect reproductive success of the plant? We observed the lowest fruit set in the microbial inoculated treatment, which recorded the highest abundance of Cluster 6. Besides, we found significant negative associations between the proportion of the ASVs of Cluster 6 and fruit set based on the GLMM analysis, while we did not find such associations for seed set. Since *Erwinia* and *Pseudomonas* include plant pathogens that trigger abortion of flowers and fruits (Goumas et al. 1999; Llontop et al. 2020; Marre et al. 2022; Young 1988), some of these bacteria might have negatively affected fruit development by infecting a reproductive organ. However, we cannot rule out other potential explanations about the correlation. Fungi, rather than bacteria, that show similar changes among the treatments with Cluster 6 might have been the cause of the low fruit set. Alternatively, bacteria and/or fungi dominant on old flowers might have indirectly decreased fruit set by changing the microenvironment of flowers. Some flower microbes are known to inhibit pollen germination or pollen tube growth (Christensen et al. 2021; Eisikowitch et al. 1990), while the effects of microbes on other post-pollination processes are largely understudied (Cullen et al. 2021). On the other hand, it is unlikely that pollinator behavioral response affected the fruit set as reported in previous studies (Sobhy et al. 2018; Vannette et al. 2013), because we hand-pollinated the flowers under the fruit set monitoring.

## 5. CONCLUSION

Compared with other floral traits, lifespan of flowers has received surprisingly little attention. Some previous studies elegantly explained variation among species based on the balance of the production and maintenance cost and benefit in terms of reproductive outputs (Ashman 2004). However, it has rarely been argued why plants so quickly replace cheap, fragile flowers during the flowering period rather than maintaining expensive but durable flowers. Short lifespan of the flower may be a defense measure of the plant to escape from negative effects of the microbes that proliferate on flowers during anthesis, though these microbes may not cause apparent symptoms. It is increasingly recognized that flowers have characteristic microbial communities, but the effects and implications of these communities have just started to be explored and our knowledge about them is still very fragmentary (Cullen et al. 2021; Rebolleda-Gómez et al. 2019; Rowe et al. 2020; Vannette 2020). Future studies will elucidate the impacts of microbes on plant reproductive ecology deeply embedded in the evolution of angiosperms (Rebolleda-Gómez et al. 2019).

## Acknowledgements

We are grateful to Otsu city for permission to conduct field work in Seta Park. This research was supported by JSPS KAKENHI Grant Numbers 19K22455 and 20H03324.

## Author contributions

NJE, MU and SS designed the study. NJE conducted field observation, sampling and amplicon sequencing with the assistance of SS and MU, respectively. SS and NJE wrote the manuscript. All authors gave final approval for publication.

## Data availability statement

Original datasets of amplicon sequencing were deposited in DDBJ (Accession No. DRA012415).

---

**TABLE S1.** The number of the samples used to monitor fruit set for the five treatments and to analyze microbial communities.

| Treatments | Fruit set and seed production |  |  |  |  | Microbial community |  |  |  |  |
| --- | --- | --- | --- | --- | --- | --- | --- | --- | --- | --- |
|  | N1 | N2 | N4 | N5 | N6 | N1 | N2 | N4 | N5 | N6 |
| Bud | - | - | - | - | - | 5 | 4 | 4 | 5 | 4 |
| Open | 10 | 26 | 14 | 16 | 19 | 5 | 5 | 5 | 5 | 5 |
| Net bag | 9 | 17 | 15 | 17 | 10 | 5 | 4 | 4 | 4 | 4 |
| Paper bag | 7 | 9 | 5 | 7 | 9 | 5 | 5 | 2 | 5 | 5 |
| Microbe | 6 | 11 | 5 | 7 | 5 | 3 | 5 | 4 | 5 | 4 |
| Water | 4 | 3 | 6 | 7 | 7 | 5 | 4 | 4 | 5 | 5 |

**TABLE S2.**
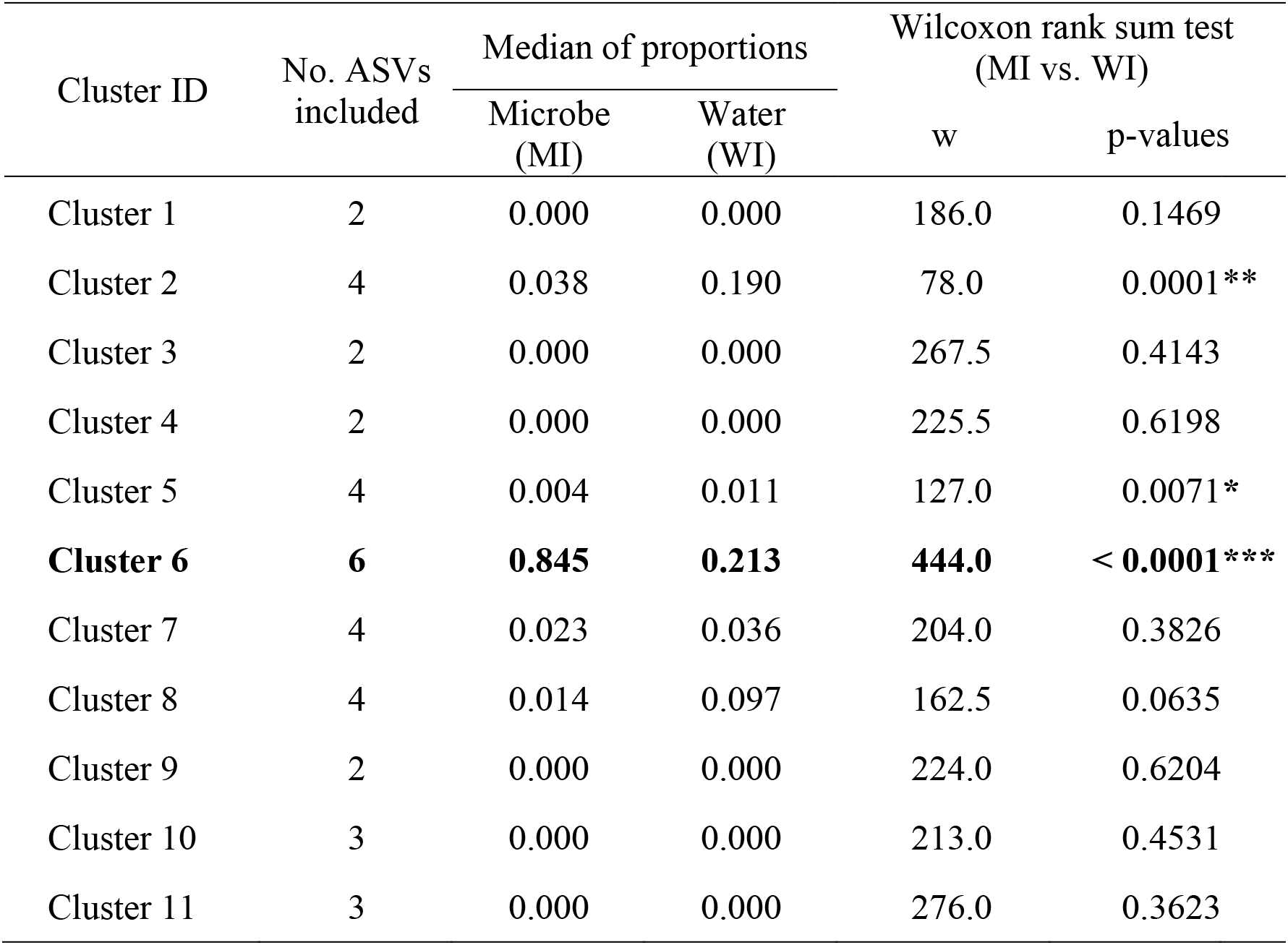
Characteristics of the 11 clusters. The number of ASVs included, the median proportion in the samples of the microbial inoculated (MI) and water sprayed (WI) treatments, and p values of Wilcoxon rank sum tests in the comparisons of the proportions between MI and WI are shown. The asterisks indicate that the proportions are significantly different between the treatments (*, p < 0.01;**, p < 0.001; ***, p < 0.0001). Cluster 6, which shows significantly higher proportions in MI than in WI, is highlighted with boldface type.

**FIGURE S1.**
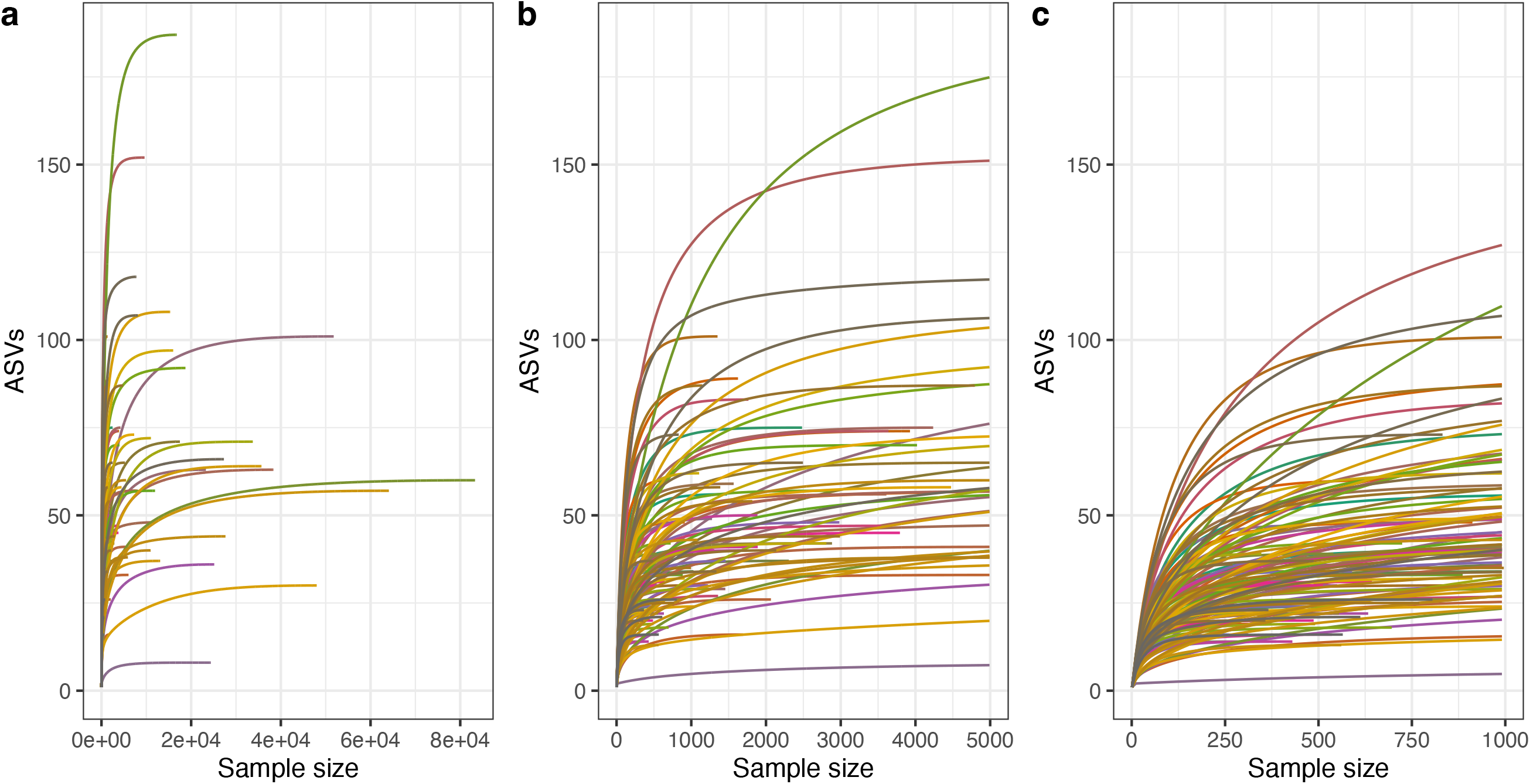
Rarefaction curves of the samples. The vertical axis shows the number of observed prokaryotic ASVs. The number of sequences per sample is shown on the horizontal axis. The three panels show the same graph with different ranges of the horizontal axis.

**FIGURE S2.**
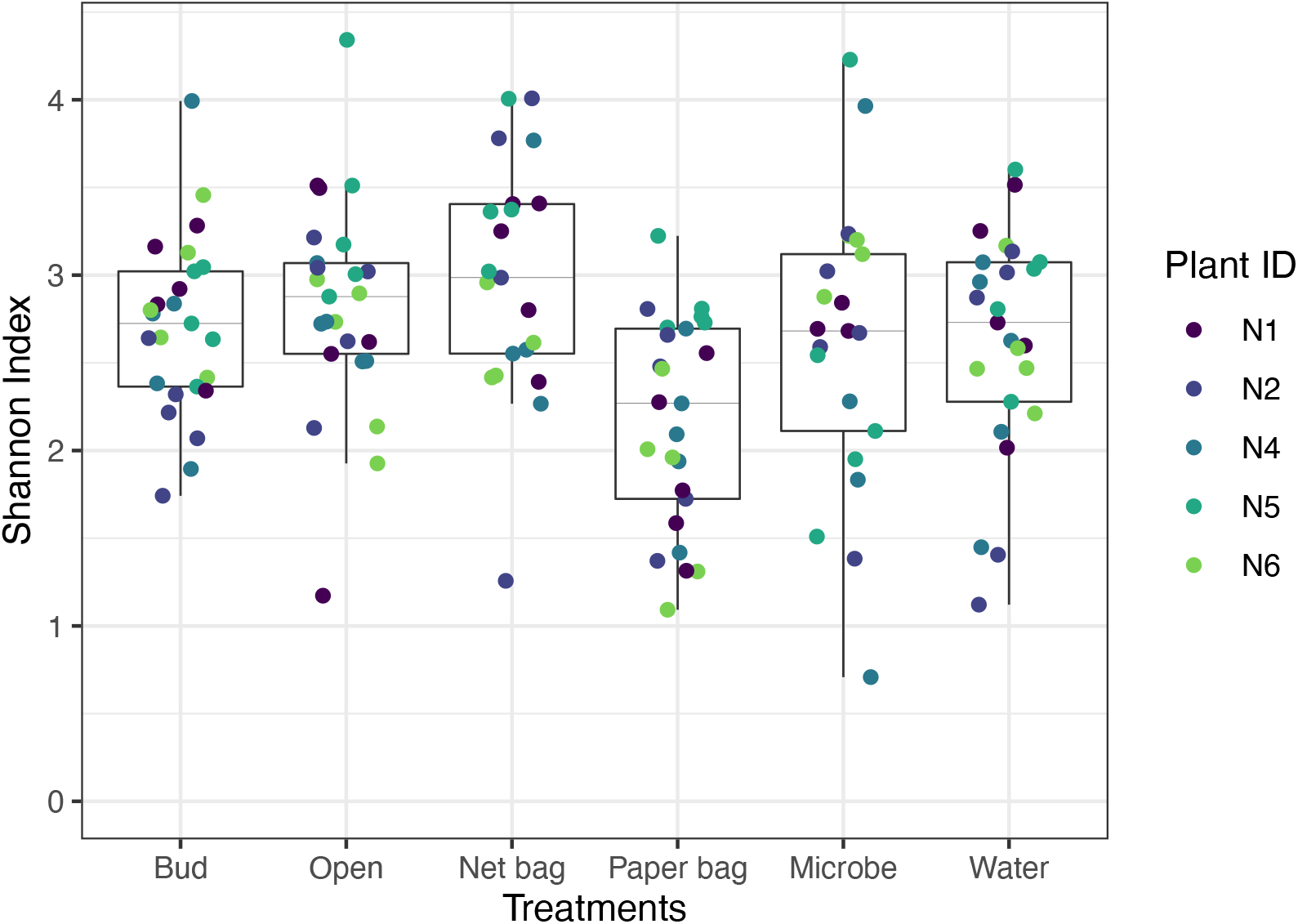
Variation in the prokaryotic diversity among the treatments and samples. The diversity was significantly different among the treatments (GLMM, χ^2^ = 22.234, p = 0.0005). Colors indicate the five individual plants.

**FIGURE S3.**
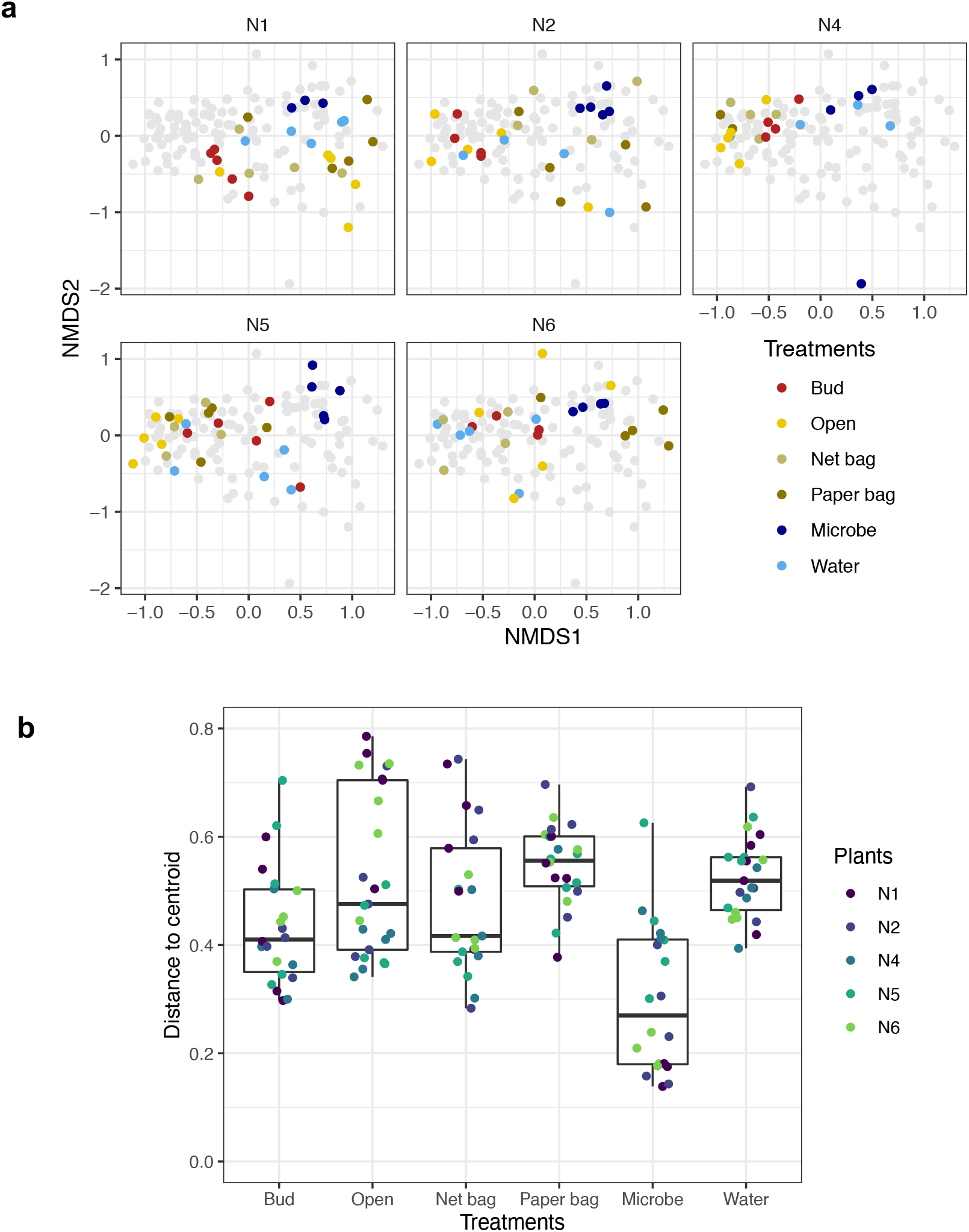
Variation of the prokaryotic communities on the flowers under different treatments. (a) NMDS plots of the microbial communities of flower buds and flowers under different treatments. In the five panels, all microbial samples are plotted with grey circles, and samples from each plant are overlayed individually. The five treatments are indicated by different colors. (b) Distribution of the distances from the treatment centroid. Variation in the microbial communities within a treatment was significantly different among the treatments (ANOVA, df = 5, F = 8.60, p < 0.0001). Colors indicate the five individual plants.

**FIGURE S4.**
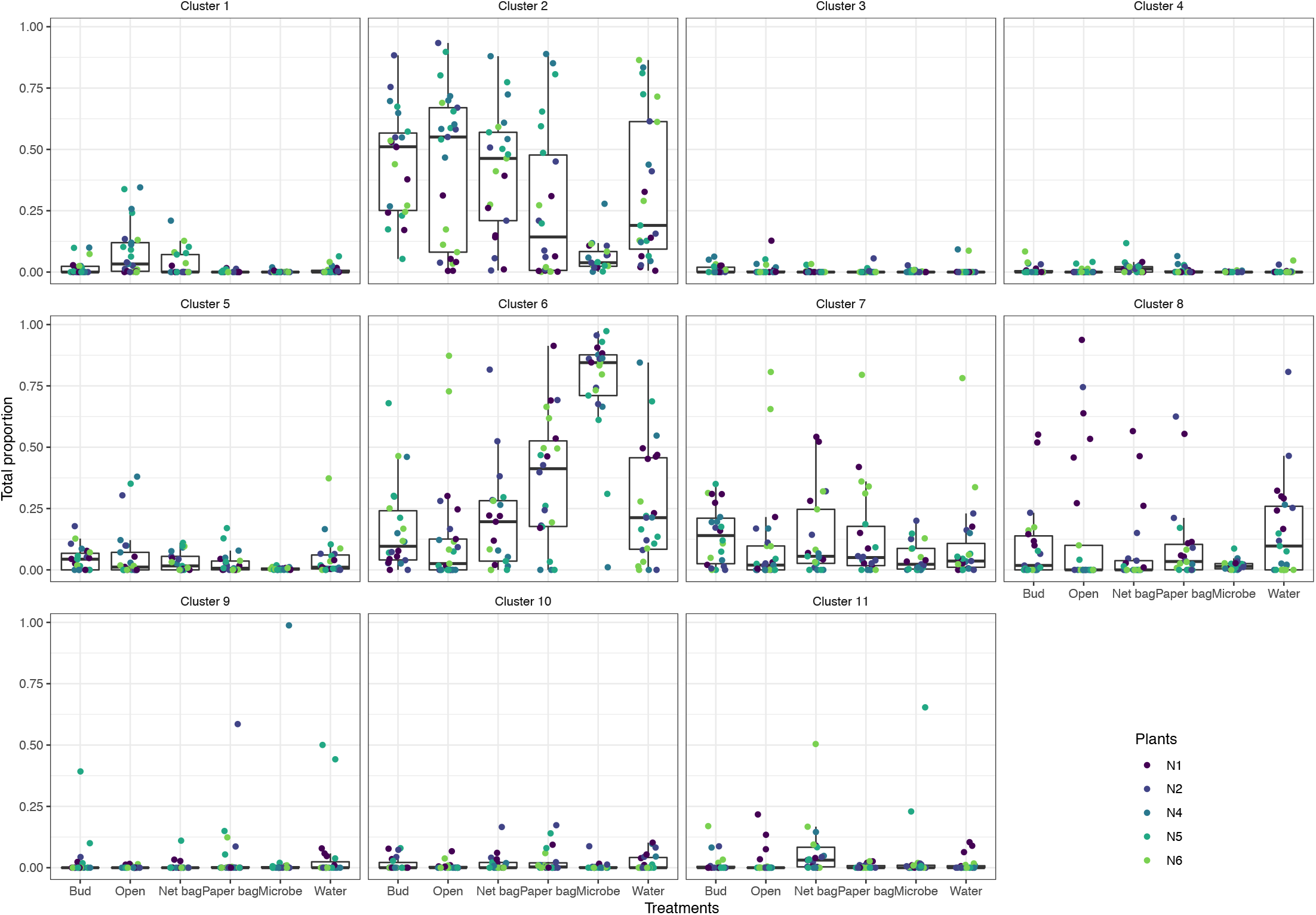
Variation in the total proportions of the 10 ASV clusters among the flower bud and flowers under the five treatments. Colors indicate the five individual plants.

